# Methylparaben dampens virulence and transmissibility of the *Drosophila* pathogen *Pseudomonas entomophila*

**DOI:** 10.1101/2024.10.11.617782

**Authors:** Youn Henry, Berta Canal-Domènech, Christine La Mendola, Tadeusz J. Kawecki

## Abstract

In the last 20 years, *Pseudomonas entomophila* (Pe) has emerged as a model to explore insect immunity to bacterial intestinal pathogens. Laboratory studies evidenced multiple detrimental effects of Pe on *Drosophila melanogaster*. However, these effects require that the bacteria are ingested in extremely high concentrations of 10^10^ – 10^11^ CFU per mL (OD_600_ 20 – 200), questioning the relevance of this pathogen in nature. Here, we tested whether the need for such high doses may be due to protective effects of the food preservative methylparaben (Nipagin), a standard ingredient of artificial *Drosophila* diets. While significant mortality in flies fed diet containing standard methylparaben concentration required doses of >10^10^ CFU per mL, when methylparaben was absent we could observe mortality using 500,000× lower doses. Consistent with these results, we demonstrated strong bactericidal properties of methylparaben on Pe *in vitro*. In the absence of methylparaben even the smallest inocula (10^5^ CFU per mL) led to high bacterial loads (10^6^ CFU per fly) after several days, indicating the ability of Pe to grow and overcome the flies’ defenses. We also demonstrate that in the absence of methylparaben, infected flies could easily transmit the pathogen to other adults and to offspring, resulting in high mortality and thus highlighting the potential of Pe as a pathogen of *Drosophila* in nature. Our study also underscores that careful consideration should be given to food additives used in standard diets in laboratory research on host-pathogen interaction.

**Importance:** Accurate characterization of pathogen infections requires appropriate experimental methodologies. Infections of insects with Pe are frequently studied using fruit flies as a model organism, with laboratory cultures typically maintained on artificial media containing various food preservatives. In this study, we show that one commonly used preservative, methylparaben, significantly influences the outcome of oral infections with Pe. We found that minimal infection doses, far below the standards of the field, could be still lethal to flies raised on media without methylparaben. This increased virulence was also associated with increased transmission of the pathogen, both from infected adult flies to their offspring and to uninfected adults. Our findings show how subtle variations in experimental conditions can profoundly affect how we perceive pathogenic threats.

## Introduction

Since Vodovar et al. (2005) first described *Pseudomonas entomophila* (Pe) and characterized its pathogenicity for fruit flies, this Gram-negative Gammaproteobacterium has been widely used as a model in fields such as immunity, ecology, evolution or sexual selection (Chakrabarti et al., 2012; Dieppois et al., 2015; Faria et al., 2015; Joye and Kawecki, 2019; Kuraishi et al., 2011). Although Pe can infect multiple insect species, most of the research focused on its interaction with the fruit fly *Drosophila melanogaster*. When ingested in sufficiently high doses, the toxins released by this pathogen in conjunction with reactive oxygen species (ROS) produced by the fly itself as part of its immune response leads to the rupture of the gut epithelium and eventually to death of the flies (Chakrabarti et al., 2012; Opota et al., 2011; Prakash et al., 2023; Vodovar et al., 2005).

One peculiar feature of most studies that performed oral infections of Pe is the dose of pathogen used (see table S1 for an overview of doses used in different studies). Following the example of the first publication on the topic (Vodovar et al., 2005), researchers usually feed the flies highly concentrated Pe suspensions (Optical Density at 600 nm 20–200), resulting ingested loads of 10^5^-10^8^ CFU per fly within a day (Gupta et al., 2013; Opota et al., 2011; Siva-Jothy et al., 2018). This practice does not reflect any ecologically realistic situation as such concentrations can only be obtained artificially, by centrifugation; Pe is unable to grow beyond OD_600_ ≈ 3 even under ideal laboratory conditions (personal observation). The need to use high oral doses is even more surprising when compared to the low doses used for systemic infections by pricking with a needle, which are lethal even with inocula of *c*.*a*. 50 bacteria per fly (Prakash et al., 2023). The protective function of the epithelial gut barrier in conjunction with the peritrophic matrix are often the main arguments used to explain the discrepancy in doses for oral vs systemic infections (Hegan et al., 2007; Martins et al., 2013; Nehme et al., 2007). Pathogen clearance in the gut has been reported to occur in less than 16h, and attributed to an efficient immune system (Bou Sleiman et al., 2015; Opota et al., 2011; Vodovar et al., 2005). However, this explanation may not be entirely satisfying given the magnitude of the difference in the lethal dose between oral and pricking infections.

All published experimental studies working with the Pe-*Drosophila* system have been performed in the laboratory, with flies and larvae fed artificial diet media composed of agar, nutrients (yeast, sugar, often cornmeal or another source of starch) and antifungal preservatives. These preservatives, such as propionic acid but primarily methylparaben (CAS No. 99-76-3, hereafter named “mp” and alternatively named Nipagin, Tegosept, or Moldex in the literature), attracted our attention. Although researchers use them to avoid mold in batch fly cultures, potential side effects for flies or for fly-associated bacteria have been largely disregarded. While antibacterial properties of mp have been extensively described decades ago (e.g. Aalto et al., 1953; Close and Neilsen, 1976; Crovetto et al., 2017; Soni et al., 2002), it is only recently that several studies explicitly pointed out the important effects of mp on growth of fly microbiota (Keebaugh et al., 2019; Obadia et al., 2018; Téfit et al., 2018). In particular, mp showed marked growth inhibition on *Acetobacter* and to a lesser extent on *Lactiplantibacillus*, two abundant commensal genera in fruit flies. In fact, the use of mp has been a central point of debate in the case of a failed replication study looking at microbiota effect on fly behavior (Leftwich et al., 2017; Leftwich et al., 2018; Rosenberg et al., 2018; Sharon et al., 2010). Finally, we noted that the only two studies which employed low Pe infection doses were also the ones not using any antifungal preservative (Beebe et al., 2015; Hegan et al., 2007) (table S1). All these elements point to the hypothesis that mp also exerts negative effects on Pe, thereby explaining the unrealistically high pathogen doses required to harm flies.

In this study, we first tested whether mp (as well as propionic acid, the second preservative often used in Drosophila media), affected Pe growth *in vitro*. Then, to assess the protective effect of mp on flies exposed to Pe, we subjected flies maintained on diet with and without mp to oral infection at various doses and quantified their survival and Pe load. The results demonstrated that in the absence of antimicrobial agents, Pe is highly virulent at doses six orders of magnitude lower than those typically used in experiments. Therefore, we also investigated the potential for adult-to-offspring and indirect adult-to-adult transmission, exploring the consequences of our findings on the ecology of this insect-pathogen relationship.

## Results

### Methylparaben and propionic acid are bactericidal to Pe *in vitro*

We verified that mp is harmful to Pe *in vitro*, at a concentration typical of what is used in fly cultures (figure 1, table S2). While Pe grew about 10,000-fold over 24 h in liquid culture in the absence of mp, we could not recover any colonies after a 24h incubation in growth medium containing 0.2% mp (Δ CFU _(control – *mp*)_ at t+24h = −3.9e+08 [−7.1e+08; −8.7e+07]). This demonstrates a bactericidal effect of mp on Pe, although we noted that short exposures of 6h were only inhibiting growth without killing cells, indicating a more bacteriostatic effect on short-term (personal observation). Similarly to mp, 0.5% propionic acid also showed strong bactericidal properties on Pe (figure S1)

**Figure 1.**
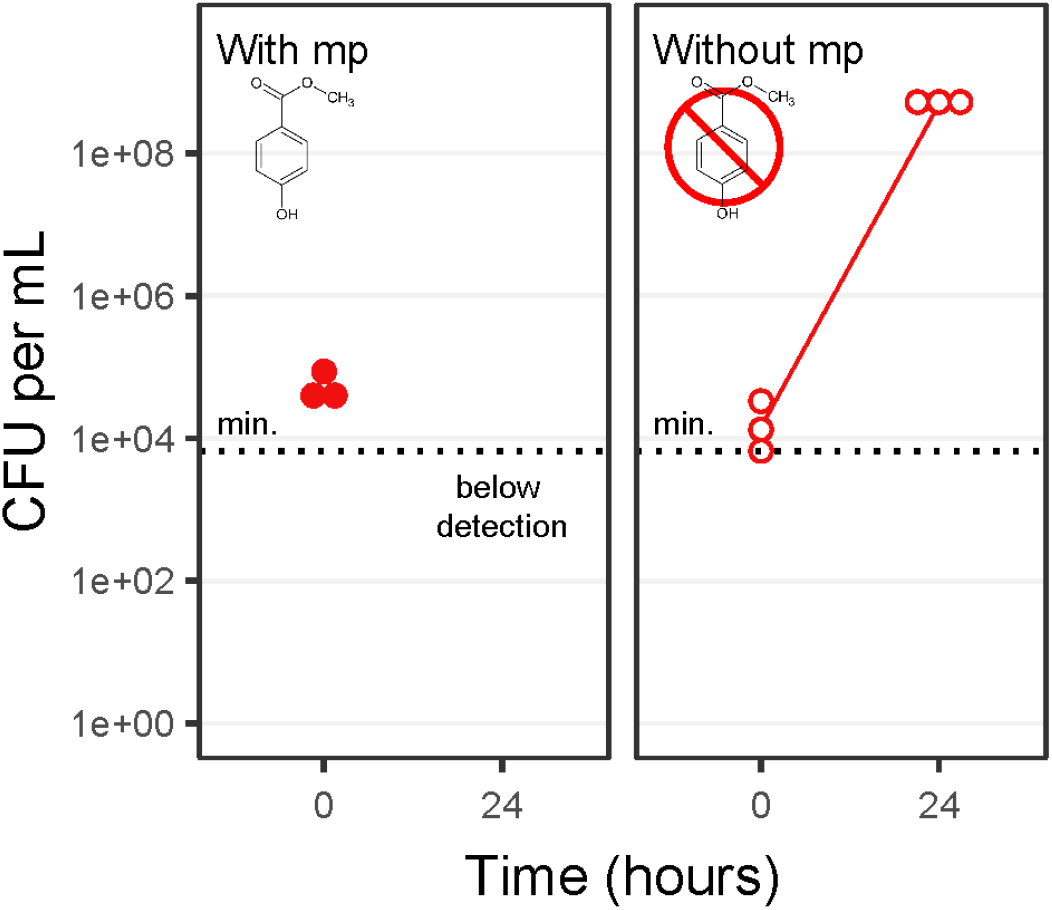
Effect of mp on Pe growth in vitro. Dots represent raw CFU measurements of N = 3 replicates. The solid dots are the condition with mp, and the open dots with dashed line are the condition without mp. The lower limit of detection was 6.6×10^3^ CFU per mL (indicated with the horizontal dotted line), and the higher limit of detection was 1.0×10^10^ CFU per mL.

### Flies die from low-dose Pe infections in the absence of methylparaben

Then, we explored the consequences of dietary mp for the outcome of oral Pe infection in adult flies. In the presence of mp, flies infected with various doses of Pe showed survival curves matching the expectations (figure 2A, top row). Only the highest dose (OD_600_ 50) led to a severe mortality within a week of infection (Δ survival _(control – OD *50*)_ at final time = −0.83 [−0.71; −0.94]). Almost no deaths were observed among flies infected with lower doses in the OD_600_ 0.0001 – 1 range (all Δ survival overlap 0, see table S3). In contrast, all pathogen doses we used were highly lethal for flies maintained on food without mp (table S3). Even the smallest concentration (OD_600_ 0.0001) led to significant mortality after a week (Δ survival _(control – OD *0*.*0001*)_ at the final timepoint = −0.38 [−0.19; −0.55]), although not as strikingly as the larger doses.

**Figure 2.**
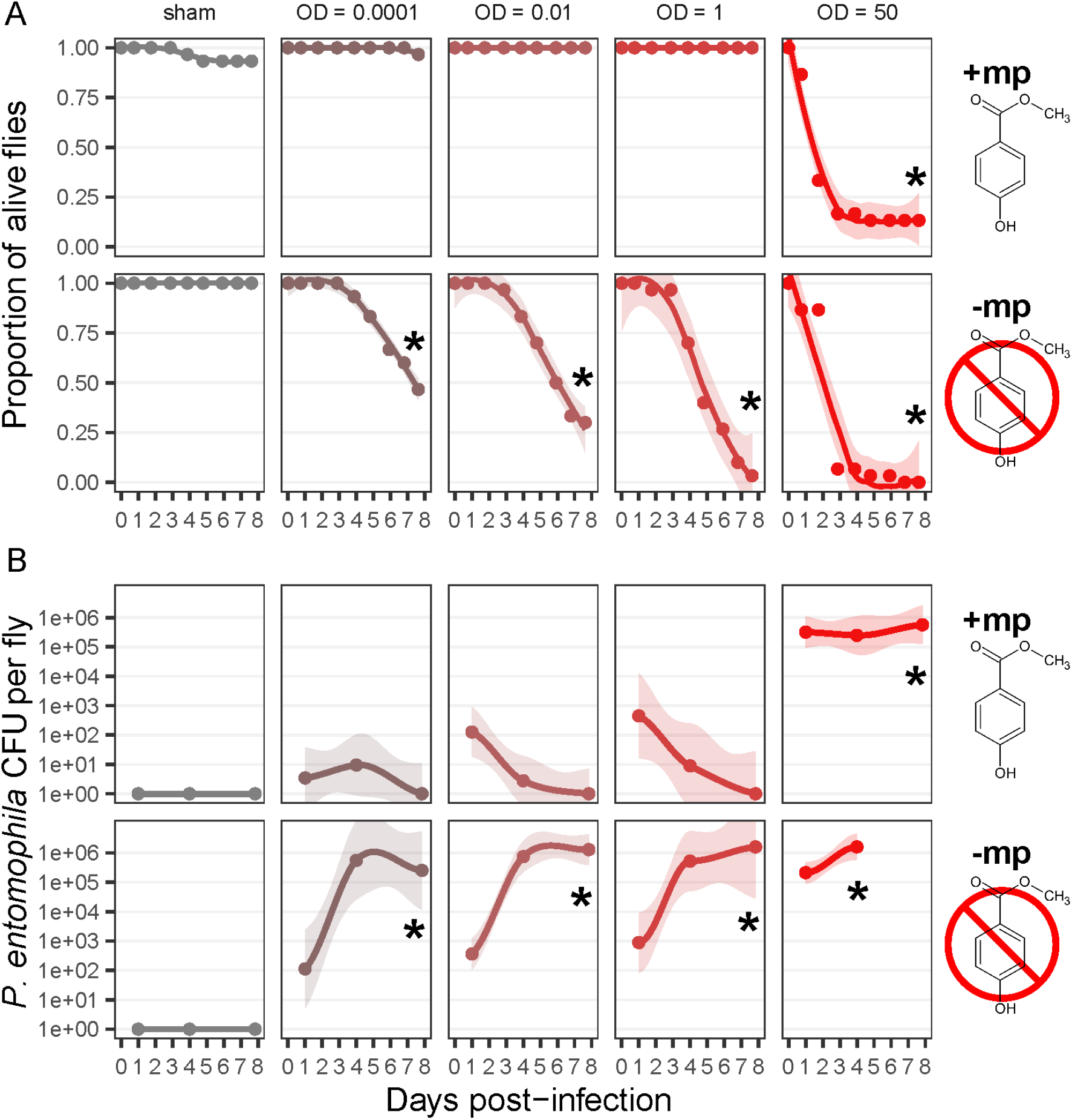
Survival (A) and pathogen load (B) of female flies exposed to different doses of pathogen (in columns), and in the presence or absence of mp in the diet (in lines). The color gradient represents the increasing Pe dose, from grey (sham infection) to red (highest infection dose of OD_600_ 50). In (A), each dot is the average survival proportion of N = 2 replicates, with 15 flies each. The line represents a loess regression on non-averaged proportions and the shaded ribbon the 95% confidence interval on this regression. The “*” indicates non-overlapping credible intervals at final time from the posterior distribution, compared to the sham-infected condition. In (B), each dot represents the average number of Pe CFU per fly, out of two flies sampled from N = 3 vials. The line represents a loess regression on non-averaged CFU and the shaded ribbon the 95% confidence interval on this regression. The limit of detection range was 40 – 3.2×10^6^ CFU per fly. Missing points are cases where all flies died before the measurement.

**Figure 3.**
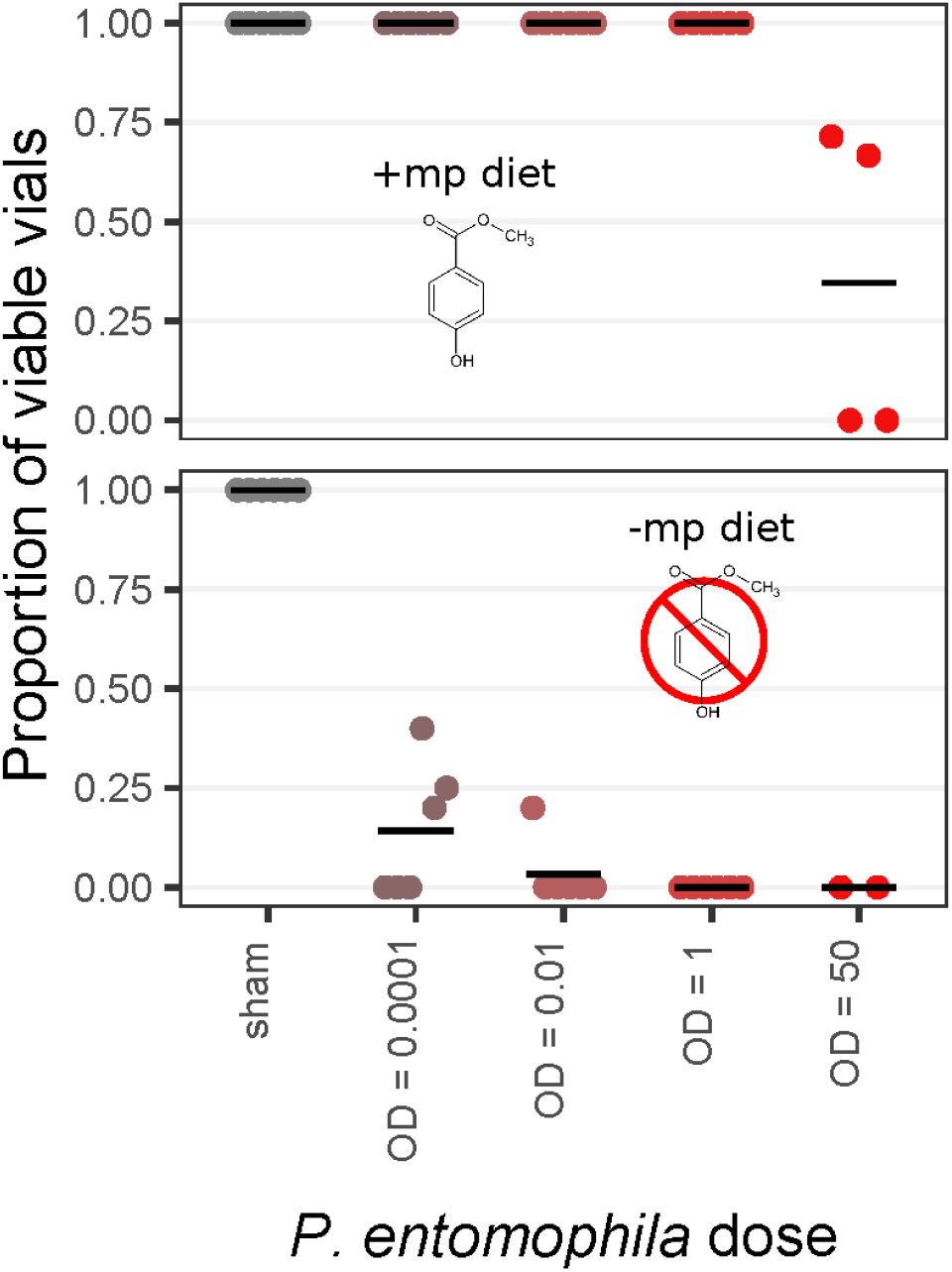
Proportion of viable offspring vials depending on the presence or absence of mp in the food. The color gradient represents the increasing Pe dose, from grey (sham infection) to red (highest infection dose of OD_600_ 50). Each dot represents the proportion of viable vials at one vial change timepoint, for N =6 timepoints. The proportion was calculated as the number of viable vials out of the total number of considered vials for a given timepoint and treatment (5 vials when all considered). Vials were considered viable if more than 5 pupae and less than 5 dead larvae were visible; vials were considered not viable if less than 5 pupae and more than 5 dead larvae were visible; other vials were not considered for the analysis.

The pathogen load well matched the survival data, with flies showing noticeable mortality only when their Pe load was high (figure 2B). In the presence of mp, Pe CFU counts at final time were different from the sham control only at OD_600_ 50 (Δ CFU _(control – OD *50*)_ at final time = 4.2e+05 [3.7e+02; 1.5e+06]), but in the absence of mp all infection doses resulted in high loads of 10^5^-10^6^ CFU per fly (table S4A).

### Daily transfer improves fly survival to Pe

The above results imply that, in the absence of mp, Pe multiplies even if initially at very low dose. This multiplication may be happening both in the fly gut and in the fly food medium. To test for the importance of Pe multiplication in the fly food medium we tested the consequences of a daily transfer of infected flies to new vials with fresh food, thus only allowing Pe present in fly guts and on their body surface to persist through the transfers. This daily change for fresh vial rescued flies’ survival, even in the absence of mp (figure S2). By resetting Pe growth every day and preventing it from thriving on fly food, we managed to mitigate Pe-induced mortality at lower doses (Δ survival _(control – OD 0.0001)_ and Δ survival _(control – OD *0*.*01*)_ both overlap 0; see table S3), but observed a mild mortality at OD_600_ 1 (Δ survival _(control – OD *1*)_ = −0.16 [−0.04; −0.28]) and a strong mortality only at OD_600_ 50 (Δ survival _(control – OD *50*)_ = −0.94 [−0.88; −0.99]). Although average CFU quantification visually showed a two to three orders of magnitude reduction in load with transfer, we could not confirm any reduction in individual contrasts with our model (all Δ CFU _(transfer – no transfer)_ overlap 0, see table S4B). Overall, these results suggest that the lethal effects of Pe are magnified by reinfection with bacteria that multiply in the food medium.

### High transmissibility of Pe in the absence of methylparaben

By keeping the vials of the daily transfer condition, we could determine the pathogen transmission to individuals of the next generation by scoring the egg-to-pupae viability (figure 3). Both the presence of mp and the Pe dose applied to parents affected the egg-to-pupae viability (*X*^2^_(df = 1, N=6)_ = 24.6, *p* < 0.001 and *X*^2^_(df = 1, N=6)_ = 22.7, *p* < 0.001 respectively). In the presence of mp, all vials had viable pupae except in the OD_600_ 50 condition which showed significant but variable mortality. In the absence of mp, all parental Pe doses led to the almost complete extinction of the next generation.

In the last experiment, we tested the indirect transmission and pathogenicity of Pe. We observed that 24h were enough for a single male infected with a low Pe dose (OD_600_ 0.01) to contaminate their environment and trigger a lethal infection in around 40% of the newly arrived females within a week of exposure (figure 4A; *X*^2^_(df = 1, *N* = 80)_ = 9.8, *p* = 0.001). The Pe load of these secondarily infected flies was highly variable, with some completely uninfected, and some with infections over 10^7^ CFU per fly (figure 4B). However, this variation in the infection status was well correlated to the survival within a given vial, showing that flies died more in vials with larger Pe abundance (figure 4C).

**Figure 4.**
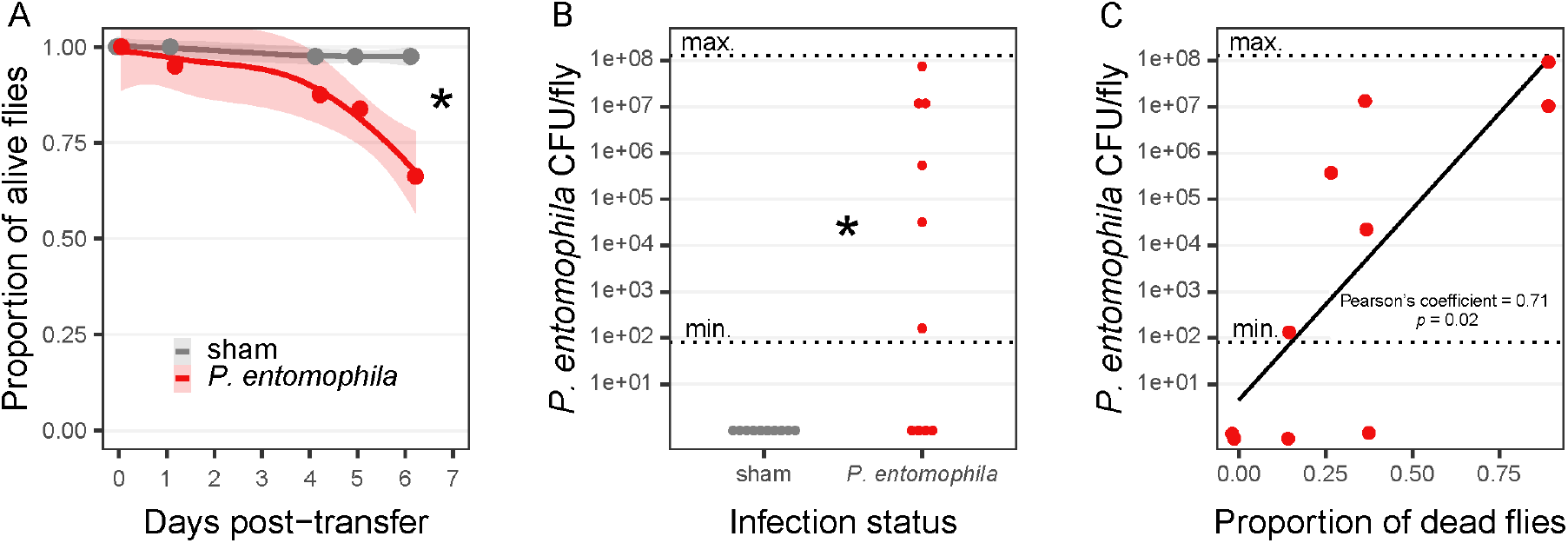
Survival (A), Pe load (B), and correlation between survival and Pe load (C), in the case of an indirect adult-to-adult transmission of Pe infection. Single sham-infected or Pe-infected males were transferred in a vial containing −mp diet, removed after 24h, and replaced with 8 females on which we measured survival and Pe load (see methods for details). In (A), each dot is the average survival proportion of N = 10 replicates, with 8 females each. The line represents a loess regression on non-averaged proportions and the shaded ribbon the 95% confidence interval on this regression. The “*” indicates a significant survival difference at final time (*X*^2^_(df = 1, *N* = 80)_ = 9.8, *p* = 0.001). In (B), each dot is the CFU count at final time (6 days) of a single fly sampled from a single replicate (N = 10 replicates). The “*” indicates a significant CFU difference at final time (Mann-Whitney U, *p* = 0.005). In (C), dots combine the proportion of dead flies in a vial (x axis) with the Pe load at final time from a fly sampled in the same vial (y axis), only keeping the Pe-infected condition. For (B) and (C), the limit of detection range, indicated with the horizontal dotted lines, was 80 – 1.2×10^8^ CFU per fly. For all plots, colors represent the infection treatment of the males that preceded the females in the vials, with gray indicating sham-infected males, and red Pe-infected males with OD_600_ 0.01.

## Discussion

In this study, we tested the effect of methylparaben (mp), a commonly used antifungal preservative in artificial *Drosophila* diet, on the pathogenicity of the model pathogen *Pseudomonas entomophila* (Pe). We observed strong antimicrobial properties of mp against Pe in both *in vitro* and *in vivo* experiments. Specifically, when all preservatives were removed from the diet, Pe was highly lethal at doses six orders of magnitude lower than those typically used in literature (e.g. Vodovar et al., 2005). This finding likely explains why earlier research often relied on extremely high doses of *Pe* to investigate gut infections (table S1). In those cases, due to the protective action of mp, flies were likely exposed to weakened or dead pathogen cells which only posed a significant threat in large quantities. This conclusion is also supported by the observation of strong mortality in flies chronically exposed to heat-killed Pe (Paulo et al., 2024).

In line with this interpretation, we show that the lethality was directly associated with the gut load of alive Pe cells, and only appeared when the bacterial load passed a certain threshold at *ca*. 10^5^ CFU per fly. However, our experimental design could not establish a directional causal link between survival and Pe load. It is possible that individuals on the verge of dying could not maintain gut homeostasis, allowing for Pe to grow out of control, or conversely that a critical mass of pathogen was required to reach a lethal tipping point (Duneau et al., 2017; Opota et al., 2011; Vijendravarma et al., 2015).

A puzzling case is the stability of the pathogen load in flies infected with the highest Pe dose on +mp diet. Despite the presence of mp, the pathogen load was maintained for a week at 10^6^ CFU per fly, which is probably close to the physical capacity of the fly gut. We suggest that this stable load might be a manifestation of the ‘inoculum effect’, a phenomenon of antibiotic resistance appearing when exposing massive cell densities to antibiotics (Brook, 1989). While this phenomenon is easily understood when bacteria can degrade the antibiotic compound (*e*.*g*. β-lactamases for β-lactam antibiotics), other mechanisms that could be at play with antimicrobial such as mp have not been clearly established (Lenhard and Bulman, 2019; Tan et al., 2012).

Experimental conditions without mp revealed that Pe was not only deadlier than expected at lower – and presumably more ecologically realistic – doses, but also more transmissible between individuals. More precisely, we evidenced adult-to-environment-to-adult Pe transmission, showing that no social physical interaction was required for transmission in the absence of mp. This is consistent with the prediction that horizontal transmission through the environment is better suited for highly virulent pathogens of non-eusocial insects (Antonovics et al., 2017; Cory, 2015; Fries and Camazine, 2001). Although Pe transmission success was imperfect, we still observed clear mortality that was directly correlated with Pe load at final time. The combination of low initial Pe concentration combined with the indirect transmission through a single infected individual likely created conditions conducive to demographic stochasticity, explaining the observed variability in survival and Pe load (Kennedy et al., 2014).

The transmission of Pe was not always directly threatening to adult flies. A short-term exposure to Pe had no lethal consequences for adults regularly moving to new uncontaminated food sources, as flies presumably often do in nature. It suggests that good sensory detection and behavioral avoidance of contaminated patches are profitable traits even for flies that are already infected, as evidenced previously (Babin et al., 2014). Yet, infected adults still had very low fitness because they transmitted the pathogen to their offspring. This was highly lethal to the larvae which, in contrast to adults, have a reduced ability to move to new uncontaminated food patches.

Overall, our findings give support to the idea that Pe is an ecologically relevant natural pathogen of *Drosophila* (Vodovar et al., 2005), which would be difficult to conceive based on the extreme doses used by most studies. This pathogen exhibits characteristics – specifically, low-dose virulence and high transmissibility – that make it suitable for exploring ecological questions, such as population-level bacterial dynamics in natural or semi-natural experimental setups. Additionally, while not necessarily invalidating their conclusions, our results shed new light on past studies. For example, Martins et al. (2013) showed that evolutionary trajectories of flies exposed to Pe depended on the infection route (oral or injury). It is possible that what was interpreted as a “route” effect was actually a “mp exposure” effect, as systemic infections appear unaffected by mp (e.g. Prakash et al., 2023). Consequently, rather than selecting for stronger immune responses against Pe, the evolution of resistance may have favored individuals with lower ROS production to avoid sepsis syndrome or those with decreased appetite to ingest less Pe during the infection step.

In this study, we focused mainly on Pe as the pathogen and mp as the antibacterial agent. Yet, we predict that our results could be extended to other pathogens and other preservatives. Propionic acid, another widely used preservative in artificial fly diet, was able to kill all Pe cells after 24h of exposure in vitro, similar to mp. Interestingly, the only study that used high doses for oral Pe infections without mp included propionic acid as a substitute (Deshpande et al., 2022). It reinforced our idea that multiple preservatives could lead to similar pathogen protection. Being broad-spectrum antimicrobial agents, food preservatives could affect other species besides Pe. Indeed, inhibitory effects of mp have been observed in several pathogen species (Aalto et al., 1953; Close and Neilsen, 1976; Tattawasart et al., 1999), and both *Pseudomonas aeruginosa* and *Serratia marcescens* require large oral infections to kill flies maintained on diet containing preservatives (Fink et al., 2016; Mulcahy et al., 2011).

To conclude, we demonstrate that the oral pathogenicity of Pe has been historically underestimated due to the common presence of mp in artificial fly diets, a finding that could be relevant for other pathogens and other preservatives as well. Nevertheless, simply removing antimicrobial preservatives in future studies may not fully resolve this issue. Pathogens could proliferate outside their hosts in preservative-free environments, as the insect exploration behavior seeds the medium with bacteria. This could result in situations where the environmental load of the pathogen eventually builds up to lethal levels, regardless of the initial dose, as observed in this study. Ideally, studies of Pe should prevent environmental growth while still using preservative-free diets. One approach to achieve this is by performing repeated transfers onto fresh diet, which efficiently limits bacterial growth. This method was employed successfully in this study and in previous work (Blum et al., 2013). Our findings demonstrate that Pe is capable of behaving as a true fly pathogen and transmitting between individuals in scenarios beyond artificial extreme infections. This insight encourages ecology-oriented studies investigating Pe dynamics in fly populations, or evolution of virulence through generations. By adopting more accurate methodologies, we can gain a deeper understanding of pathogen-host interactions and improve the reproducibility and relevance of experimental results in the field of fly immunity.

## Methods

### Fly maintenance

For all experiments, we used a *Wolbachia*-free wildtype fly population collected in 2007 in the Valais, Switzerland. Stocks were kept in outbred conditions in a thermoregulated room set at 25°C ±0.5°C, 60% RH, and 12L:12D light cycle. The standard diet of flies was composed of brewer’s yeast (2% weight/volume), cornmeal (5.2% w/v), sucrose (11% w/v), agar (0.8% w/v), and methylparaben (ref H5501, Sigma-Aldrich, USA) (1.1% v/v of 20% stock solution in pure ethanol) mixed in water. The concentration of mp was in line with standard food recipes found elsewhere (*e*.*g*. see references from table S1). In the manuscript, “+mp diet” refers to the standard diet, and “-mp diet” refers to a modified standard diet without mp. We performed all fly transfers with brief CO_2_ anesthesia.

### Bacterial cultures

For all experiments, we used the L48^T^ strain of Pe (kindly shared by Bruno Lemaitre), grown overnight in LB medium (ref 240230, BD Difco, USA) at 30°C 150 rpm. When required, we standardized the concentration to the desired OD_600_ by pelleting the cultures at 3000 rpm for 5 min, discarding the supernatant, and resuspending in a sterile solution of 5% sucrose for infections or sterile 25% glycerol solution in PBS buffer for plating and storage. For each experiment, we plated Pe to verify that OD_600_ 1 corresponded approximately to 10^9^ CFU/mL.

### Effect of methylparaben and propionic acid on Pe *in vitro*

We designed a test to characterize the bactericidal or bacteriostatic effect of methylparaben on Pe *in vitro*, at a concentration comparable to what is used in artificial fly diets. In a 96 wells microplate, we added 1% of ethanol or 1% of ethanol containing 20% methylparaben to a standardized suspension of Pe (OD_600_ 0.0005) in LB medium, with 3 replicated wells per condition. We immediately plated a serial dilution of suspension from each well with PBS on *Pseudomonas* isolation agar to obtain estimates of initial bacterial concentration. We incubated the microplate at 28°C with agitation. After 24h of incubation, we plated another serial dilution with PBS from each well on *Pseudomonas* isolation agar. We counted the colonies after incubation of the plates for 20h at room temperature. We also tested the effect of propionic acid, another widely used antifungal additive, on Pe. With the same procedure and design as for mp, we compared growth of Pe exposed to 0.5% propionic acid or to 0.5% PBS buffer control.

### Oral infection procedure

We largely followed standard infection procedures, extensively described by Siva-Jothy et al. (2018). Before the infection, we starved flies for 4h in tubes containing water agar (1%), with 8-15 flies per tube. We prepared the infection vials by adding a filter paper on the surface of a new water-agar vial and pipetted down 100μL of the pathogen suspension in 5% sucrose with the desired OD_600_ or 100μL of the 5% sucrose solution only (sham control). We transferred the flies to these vials and, after 20h of infection, transferred them again to fresh vials containing food with or without mp. We then monitored the flies maintained at 25°C ±0.5°C, 60% RH, and 12L:12D light cycle, for up to eight days to record survival and Pe load. We recorded survival with daily counting of dead individuals. We measured the bacterial load in flies by plating alive individuals sampled from separate dedicated vials.

### Gut load of *Pseudomonas entomophila*

To measure the Pe load in flies’ guts, we sampled 1-2 random individuals per vial at different timepoints post-infection, depending on the experimental block. We briefly dipped the collected the flies in 70% ethanol, vortexed, discarded the ethanol, and let the flies dry for 30 min to ensure killing external bacteria. We then crushed the flies in 400μL of 1:1 mix of sterile 40% glycerol and sterile PBS, with a 2mm steel bead using a tissue homogenizer (Precellys® evolution, Bertin, France) at 4500rpm for 1 minute. After a serial dilution range, we plated 5μL of each dilution on *Pseudomonas* isolation agar medium (ref17208, Merck, Germany), and counted the colony forming units (CFU) after 20h growth at room temperature. The CFU detection range spanned from 20 – 80 CFU per fly (= 1 colony in the undiluted homogenate) up to 1.6×10^6^ – 1.2×10^8^ CFU per fly (= 200 colonies in the most diluted homogenate), depending on the experimental block.

### Effect of methylparaben and daily transfer on fly survival

Six days old individuals were sexed under CO_2_ anesthesia, keeping only females (which were likely mated, having spent several days in mixed sex groups). We then divided females into multiple vials (15 females per vial) of +mp diet or −mp diet. After 3 days on this new diet, we proceeded to infections (see infection procedure section for details). We used five concentrations of Pe (OD_600_ = 50, 1, 0.01, 0.0001 or 0) and after 20h of infection, we transferred the flies to fresh vials containing the +mp or the −mp diet. Then, we maintained the flies either in the same vial for up to seven days or transferred them daily to new vials with fresh food, recording survival rates or Pe load. This daily transfer condition allowed us to test whether Pe was stably colonizing the gut or whether it required constant replenishment *via* feeding, as observed with most bacteria from the fly microbiome (Blum et al., 2013; Pais et al., 2018). In the condition with daily transfer, we always counted the dead flies and sampled individuals for Pe load measurement just before the next transfer. Dead individuals were counted daily. They were not removed from the vial they died in, meaning they were not transferred to the new vial in the daily transfer condition. In total, we tested 20 different conditions (2 +mp/-mp diets × 2 transfer/no transfer × 5 infection doses). We used two replicate vials per condition for survival, and three replicated vials for Pe load measurement.

### Adult-offspring pathogen transmission

We kept the used vials from the daily transfer condition (see above) to assess the viability of the offspring, by counting dead larvae and pupae six days after the change (N=5 replicates per condition and per day). The variable number of individuals caused by death events strongly affected the number of eggs laid. For that reason, we opted for a binary classification of the offspring vials: a vial was considered as viable when the number of dead larvae was below five and the number of pupae exceeded five, and a vial was considered as non-viable when the number of dead larvae exceeded five and the number of pupae was below five. All vials not matching these criteria (21/300) were discarded from the analysis.

### Indirect adult-adult pathogen transmission

To investigate the extent of Pe pathogenicity at low doses and differentiate between environmental and social transmission routes, we conducted a separate experiment testing indirect pathogen transmission among adults. Four days old individuals from the stock population were sexed under CO_2_ anesthesia. We then divided individuals into multiple single-sex vials (8 females or 8 males per vial) of −mp standard diet. After 3 days on this new diet, we started the infection procedure (see infection procedure section for details). We used two infection treatments (Pe at OD_600_ 0.0001 and sham control) on male vials for 20h, and then transferred single males into fresh vials containing −mp standard diet. After 24h, we discarded the males and introduced uninfected females into each used vial. We then kept the females for six days in the same vial, recording their survival (10 replicates per condition). We also plated surviving flies at the end of the experiment to quantify Pe load.

### Statistical analysis

We analyzed most datasets in a Bayesian framework, using R (version 4.2.1, R Core Team, 2022) and the `brms` package (Bürkner, 2017) as frontends for the Stan language (Carpenter et al., 2017). We used the `tidyverse`, `tidybayes`, `bayesplot`, and `patchwork` packages for data preparation, model evaluation and plotting (Gabry et al., 2019; Kay and Mastny, 2022; Pedersen, 2022; Wickham et al., 2019).

We analyzed Pe CFU variation in the *in vitro* culture using a Bayesian linear model. Our model included mp presence and time as fixed factors, in interaction. We also included a group-level random effect associated with each replicate.

We analyzed flies’ survival after a direct Pe infection using a Bayesian binomial linear model with a logit link function, using a dataset of the survival at final time only (t = 183h). Our model included mp presence, infection dose measured as OD_600_, and vial change as fixed factors, with all interactions.

We analyzed Pe CFU load after a direct Pe infection using a Bayesian linear model, using a dataset of the CFU at final time only (mostly t = 188h, or earlier datapoints for a few vials that had no more survivors at final time). Our model included mp presence, infection dose measured as OD_600_, and daily transfer as fixed factors, with all interactions.

We fit our models using weakly informative priors inspired by McElreath (2020): Normal(0,1.5) prior for the intercept, Normal(0,1) prior for the random effect, and Normal(0,1) or Normal(0,5) priors for the fixed effects in the survival model and the CFU models respectively. We ran four chains for 10 000 iterations, with the first half of each chain used as a warmup. We give all results as the difference (Δ) of a posterior mean [95% Highest Posterior Density Intervals] compared to the control condition. By construction, our Bayesian approach did not require any correction for multiple comparison (Gelman and Tuerlinckx, 2000).

We analyzed the data from the direct and indirect Pe transmission experiments in a frequentist framework using R. For the direct transmission to the next generation, we analyzed the egg-to-pupae viability using the mixed binomial generalized linear model with a logit link function from the lme4 package (Bates et al., 2015). We included mp and Pe dose as interacting fixed effects, and the day of change as a random effect. For survival at final time, we used a mixed binomial generalized linear model with a logit link function. We included the infection status as a fixed factor and the vial identifier as an observation-level random effect to control for overdispersion (Harrison, 2015). For CFU, we compared the two distributions (infected vs sham) with a Mann-Whitney U test. Finally, we checked the correlation between survival and Pe load using Pearson’s correlation test.

## Data availability

Datasets and scripts are available on the Zenodo repository DOI 10.5281/zenodo.13837575.

## Acknowledgements

We warmly thank members of the Kawecki group who all encouraged us to pursue this work beyond the first serendipitous observations by Jaime González and YH.

**Youn Henry**: conceptualization, investigation, formal analysis, writing – original draft, writing – review & editing **Berta Canal-Domènech**: writing – review & editing **Christine La Mendola**: investigation **Tadeusz J. Kawecki**: conceptualization, writing – review & editing, supervision.

This work was supported as a part of NCCR Microbiomes, a National Centre of Competence in Research, funded by the Swiss National Science Foundation (grant number 51NF40_180575).

## Conflict of interest

Authors declare no conflict of interest.

## Notes

### Competing Interest Statement

The authors have declared no competing interest.

