## Supplementary material for "Methylparaben dampens virulence and transmissibility of the *Drosophila* pathogen *Pseudomonas entomophila*"

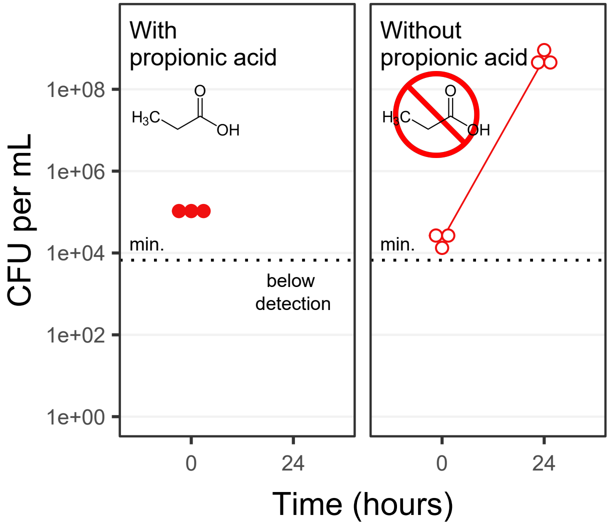

With
propionic acid
Without
propionic acid
$H_3C$
O
OH
1e+08
1e+06
1e+04
1e+02
1e+00
CFU per mL
min.
below
detection
0
24
Time (hours)

Figure S1: *In vitro* growth of Pe in the presence or not of propionic acid. Dots represent raw CFU measurements of N = 3 replicates. The solid dots with solid line are the condition with propionic acid, and the open dots with dashed line are the condition without propionic acid. The lower limit of detection was $6.6\times10^{3}$ CFU per mL (indicated with the horizontal dotted line), and the higher limit of detection was $1.0\times10^{10}$ CFU per mL.

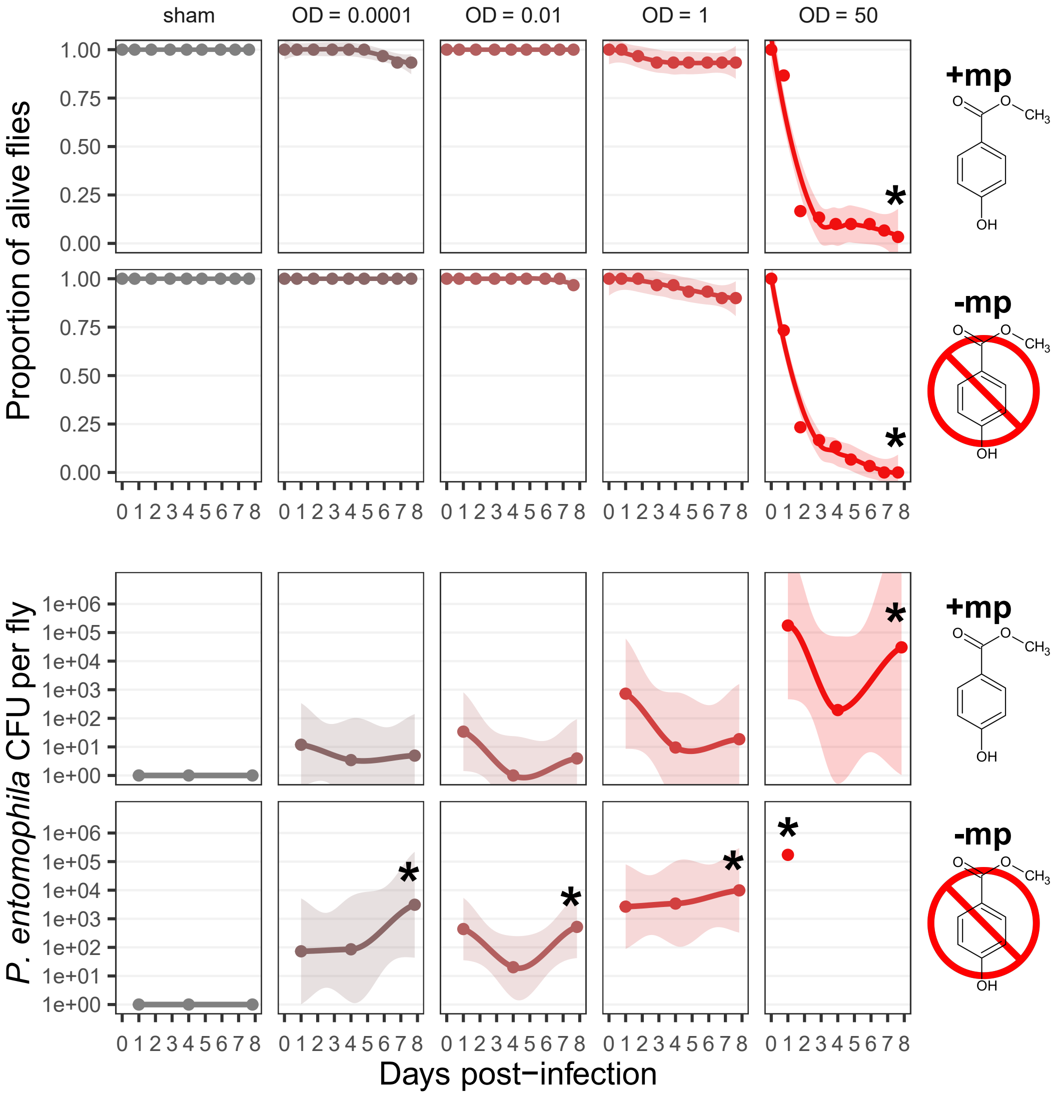

sham
OD = 0.0001
OD = 0.01
OD = 1
OD = 50
Proportion of alive flies
1.00
0.75
0.50
0.25
0.00
+mp
-mp
O
O
CH3
OH
0 1 2 3 4 5 6 7 8
P. entomophila CFU per fly
1e+06
1e+05
1e+04
1e+03
1e+02
1e+01
1e+00
*
Days post−infection

Figure S2: Survival and pathogen load of female flies daily transferred to fresh vials post-infection, exposed to different doses of pathogen (in columns), and in the presence or absence of mp in the food (in lines). The color gradient represents the increasing Pe dose, from grey (sham infection) to red (highest infection dose of $OD_{600}$ 50). For survival, each dot is the average survival proportion of N = 2 replicates, with 15 flies each. The line represents a loess regression on non-averaged proportions and the shaded ribbon the 95% confidence interval on this regression. The "*" indicates non-overlapping credible intervals at final time from the posterior distribution, compared to the sham-infected condition. For Pe load, each dot represents the average number of Pe CFU per fly, out of two flies sampled from N = 3 vials. The line represents a loess regression on non-averaged CFU and the shaded ribbon the 95% confidence interval on this regression. The limit of detection range was 40 – $3.2 \times 10^6$ CFU per fly. Missing points are cases where all flies died before the measurement.

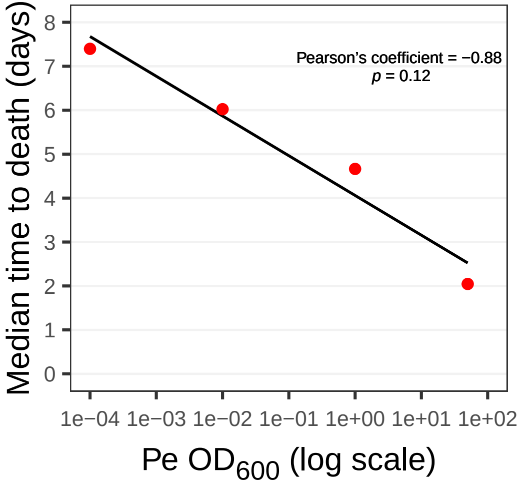
Pearson's coefficient = −0.88
p = 0.12
Median time to death (days)
0
1
2
3
4
5
6
7
8
1e−04
1e−03
1e−02
1e−01
1e+00
1e+01
1e+02
Pe OD600 (log scale)

Figure S3: Correlation between the median time to death and the Pe dose. Dots show the median time to death extracted from loess regressions of survival data (y axis) as a function of the inoculated Pe dose at the start of the experiment (x axis), only keeping the Pe-infected flies maintained on -mp diet. The black line is the linear regression on the dots.

Table S1: Summary of the literature review. Only the articles using oral Pe infections are shown here. A dash indicates missing information.

| Article | Methylparaben use | Suspected methylparaben use | Propionic acid use | Pe optical density |
|---|---|---|---|---|
| Al Zouabi et al. (2023) | - | yes | - | 200 |
| Babin et al. (2014) | yes | yes | no | 100 |
| Beebe et al. (2015) | no | no | no | 0.01-10 |
| Beehler-Evans et al. (2015) | - | yes | - | 10-20 |
| Chakrabarti et al. (2012) | - | yes | - | 100 |
| Chakrabarti et al. (2014) | yes | yes | yes | 100 |
| Deshpande et al. (2022) | no | no | yes | 200 |
| Faria et al. (2015) | - | yes | - | 50 |
| Frochaux et al. (2020) | yes | yes | yes | 100 |
| Hegan et al. (2007) | no | no | no | 1 |
| Jacob et al. (2017) | - | yes | no | - |
| Joye et al. (2019) | yes | yes | no | 100 |
| Kawecki et al. (2020) | yes | yes | no | 100 |
| Kobler et al. (2020) | yes | yes | yes | 100 |
| Kuraishi et al. (2011) | - | yes | - | 100 |
| Liehl et al. (2006) | - | yes | - | 100 |
| Loch et al. (2017) | yes | yes | no | - |
| Martins et al. (2013) | - | yes | - | 50 |
| Onuma et al. (2023) | yes | yes | yes | 200 |
| Opota et al. (2011) | - | yes | - | 100 |
| Papagiannoulis et al. (2010) | - | yes | - | 200 |
| Prakash et al. (2023) | yes | yes | no | 45 |
| Sella et al. (2024) | - | yes | - | 100 |
| Sharda et al. (2022) | yes | yes | no | 300 |
| Shibata et al. (2015) | - | yes | - | 200 |
| Siva-Jothy et al. (2018) | yes | yes | no | 100 |
| Sleiman et al. (2015) | yes | yes | yes | 100 |
| Soory et al. (2022) | - | yes | - | 200 |
| Vallet-Gely et al. (2010a) | - | yes | - | 100 |
| Vallet-Gely et al. (2010b) | - | yes | - | 100 |
| Vijendravarma et al. (2015) | yes | yes | yes | 25 |
| Vodovar et al. (2006) | - | yes | - | 200 |
| Won et al. (2023) | - | yes | - | 200 |

### Methods

We performed a simple literature review, aiming to identify the proportion of published articles using antifungal preservatives in experiments involving Pe infection of fruit flies. In Web of Science, we used the keyword combination "*Drosophila*" + "*Pseudomonas entomophila*", searching in all fields. This approach excluded some rare articles only mentioning these terms in the main text, but we deemed our research sufficient to provide a representative overview of the published record. We manually inspected the 64 initial hits and only kept articles explicitly using oral infection procedure, keeping a total of 33 articles. For all articles we collected information on the use of mp, the use of propionic acid, and the Pe $OD_{600}$ for infection. Because diet recipes were often omitted from the methods, we also recorded the "suspected use of mp". Articles with "suspected use of mp" contained no explicit information on mp use but were using some sort of "standard diet". We manually searched whether this "standard diet" contained mp in previous publications from the same research group, and if yes, we considered the article of interest had a "suspected use of mp".

### Results

We found a total of 64 hits for articles working on Drosophila infection with Pe. Out of these total hits, [33/64] articles performed oral infections, and [30/33] were likely maintaining flies on diet containing mp (down to [13/16] when only considering articles with explicit mention of the diet composition). All articles using mp performed infections with high Pe doses in the $OD_{600}$ range of 25-300. Out of the three articles with mp-free fly diet, one contained propionic acid and used $OD_{600}$ 200 for infections, and two others contained no preservative and used low infection doses ($OD_{600}$ 10, and $OD_{600}$ 1).

Table S2: Summary of model predictions for pairwise differences represented in figure 1. In the upper table, Δ are predicted for same treatment at 0 and 24h showing the time effect. In the lower table, Δ are predicted for same time with or without mp, showing the mp effect. We only interpret differences with confidence intervals not overlapping 0 as biologically meaningful and highlighted them in green or red for positive and negative differences respectively.

| Treatment | Time | Prediction | LowerCI | UpperCI |
|---|---|---|---|---|
| antifungal - control | 0 | 2.2e+04 | -1.1e+04 | 6.9e+04 |
| antifungal - control | 24 | **-3.9e+08** | -7.1e+08 | -8.7e+07 |

| Treatment | Time | Prediction | LowerCI | UpperCI |
|---|---|---|---|---|
| control | 24 - 0 | **3.9e+08** | 8.7e+07 | 7.1e+08 |
| antifungal | 24 - 0 | **-4.3e+04** | -8.0e+04 | -1.1e+04 |

Table S3: Summary of model predictions for pairwise survival differences represented in figures 2A and S2. The Δ are pairwise predicted survival differences at final time between sham infected and a given Pe infection dose, with same diet (+mp/-mp) and same vial change (with/without daily vial change). We only interpret differences with confidence intervals not overlapping 0 as biologically meaningful and highlighted them in green or red for positive and negative differences respectively.

| P.e dose | Nipagin | Vial change | Prediction | LowerCI | UpperCI |
|---|---|---|---|---|---|
| OD = 0.0001 - sham | without_nipagin | no_change | **-0.38** | -0.55 | -0.20 |
| OD = 0.01 - sham | without_nipagin | no_change | **-0.51** | -0.68 | -0.33 |
| OD = 1 - sham | without_nipagin | no_change | **-0.70** | -0.84 | -0.56 |
| OD = 50 - sham | without_nipagin | no_change | **-0.86** | -0.94 | -0.77 |
| OD = 0.0001 - sham | with_nipagin | no_change | -0.04 | -0.14 | 0.04 |
| OD = 0.01 - sham | with_nipagin | no_change | -0.01 | -0.09 | 0.06 |
| OD = 1 - sham | with_nipagin | no_change | -0.06 | -0.16 | 0.03 |
| OD = 50 - sham | with_nipagin | no_change | **-0.83** | -0.94 | -0.71 |
| OD = 0.0001 - sham | without_nipagin | daily_change | -0.05 | -0.12 | 0.01 |
| OD = 0.01 - sham | without_nipagin | daily_change | -0.06 | -0.14 | 0.01 |
| OD = 1 - sham | without_nipagin | daily_change | **-0.16** | -0.28 | -0.05 |
| OD = 50 - sham | without_nipagin | daily_change | **-0.94** | -0.99 | -0.88 |
| OD = 0.0001 - sham | with_nipagin | daily_change | -0.03 | -0.10 | 0.02 |
| OD = 0.01 - sham | with_nipagin | daily_change | 0.00 | -0.03 | 0.04 |
| OD = 1 - sham | with_nipagin | daily_change | -0.03 | -0.10 | 0.02 |
| OD = 50 - sham | with_nipagin | daily_change | **-0.94** | -0.99 | -0.88 |

Table S4: Summary of model predictions for pairwise Pe load differences represented in figures 2B and S2. In (A) the Δ are pairwise predicted Pe load differences at final time between sham infected and a given Pe infection dose, with same diet (+mp/-mp) and same vial change (with/without daily vial change). In (B) the Δ are pairwise predicted Pe load differences at final time between flies daily changed to new food or not, with same diet (+mp/-mp) and same Pe infection dose ($OD_{600}$ = 50, 1, 0.1, 0.0001, 0). We only interpret differences with confidence intervals not overlapping 0 as biologically meaningful and highlighted them in green or red for positive and negative differences respectively.

**A**

| P.e dose | Nipagin | Vial change | Prediction | LowerCI | UpperCI |
|---|---|---|---|---|---|
| OD = 0.0001 - sham | without_nipagin | no_change | **1.4e+05** | 3.6e+02 | 4.5e+05 |
| OD = 0.01 - sham | without_nipagin | no_change | **4.5e+05** | 1.1e+03 | 1.6e+06 |
| OD = 1 - sham | without_nipagin | no_change | **2.6e+05** | 7.4e+02 | 9.1e+05 |
| OD = 50 - sham | without_nipagin | no_change | **1.1e+06** | 3.8e+03 | 3.9e+06 |
| OD = 0.0001 - sham | with_nipagin | no_change | 2.5e+00 | -3.3e+00 | 1.2e+01 |
| OD = 0.01 - sham | with_nipagin | no_change | 3.1e+00 | -3.1e+00 | 1.4e+01 |
| OD = 1 - sham | with_nipagin | no_change | 2.8e+00 | -3.4e+00 | 1.3e+01 |
| OD = 50 - sham | with_nipagin | no_change | **4.2e+05** | 3.7e+02 | 1.5e+06 |
| OD = 0.0001 - sham | without_nipagin | daily_change | **5.8e+03** | 1.1e+01 | 2.0e+04 |
| OD = 0.01 - sham | without_nipagin | daily_change | **1.4e+03** | 8.8e-01 | 5.1e+03 |
| OD = 1 - sham | without_nipagin | daily_change | **1.9e+04** | 5.1e+01 | 6.7e+04 |
| OD = 50 - sham | without_nipagin | daily_change | **2.0e+05** | 4.9e+02 | 7.1e+05 |
| OD = 0.0001 - sham | with_nipagin | daily_change | 7.5e+00 | -6.6e+00 | 3.3e+01 |
| OD = 0.01 - sham | with_nipagin | daily_change | 4.7e+00 | -6.8e+00 | 2.3e+01 |
| OD = 1 - sham | with_nipagin | daily_change | 3.1e+01 | -7.4e+00 | 1.2e+02 |
| OD = 50 - sham | with_nipagin | daily_change | **1.7e+04** | 1.2e+01 | 5.9e+04 |

**B**

| P.e dose | Nipagin | Vial change | Prediction | LowerCI | UpperCI |
|---|---|---|---|---|---|
| sham | without_nipagin | daily_change - no_change | -3.5e+01 | -1.2e+02 | 5.4e+00 |
| OD = 0.0001 | without_nipagin | daily_change - no_change | -1.3e+05 | -4.7e+05 | 3.2e+04 |
| OD = 0.01 | without_nipagin | daily_change - no_change | -4.5e+05 | -1.6e+06 | 7.4e+03 |
| OD = 1 | without_nipagin | daily_change - no_change | -2.4e+05 | -9.3e+05 | 1.0e+05 |
| OD = 50 | without_nipagin | daily_change - no_change | -9.2e+05 | -4.2e+06 | 1.0e+06 |
| sham | with_nipagin | daily_change - no_change | 6.0e-01 | -2.9e+00 | 4.8e+00 |
| OD = 0.0001 | with_nipagin | daily_change - no_change | 5.6e+00 | -1.5e+01 | 3.6e+01 |
| OD = 0.01 | with_nipagin | daily_change - no_change | 2.2e+00 | -1.9e+01 | 2.6e+01 |
| OD = 1 | with_nipagin | daily_change - no_change | 2.9e+01 | -2.3e+01 | 1.2e+02 |
| OD = 50 | with_nipagin | daily_change - no_change | -4.0e+05 | -1.5e+06 | 9.2e+04 |
